# Facing the waves of negative mood: Trait mindfulness moderates the links between physiology and emotion

**DOI:** 10.64898/2026.10.30.760637

**Authors:** Ashley Michelle Campos, Gayathri Batchalli Maruthy, Genevieve Cumming, Stephanie Rodriguez, Lyndahl Himes, Bart Rypma

**Author notes:** Corresponding Author: (AMC). School of Behavioral Brain Sciences, University of Texas at Dallas. Ashley Michelle Campos: Conceptualization, Writing – original draft, Methodology, Investigation, Formal analysis, Visualization, Data curation. Gayathri Batchalli Maruthy: Conceptualization, Writing – review & editing, Supervision, Methodology, Formal analysis. Genevieve Cumming: Data curation, formal analysis. Stephanie Rodriguez: Data curation. Lyndahl Himes: Conceptualization and Funding acquisition. Bart Rypma: Writing – original draft, Supervision, Resources, Project administration, Methodology, Investigation, Funding acquisition, Conceptualization.

## Abstract

Trait mindfulness is a dispositional ability characterized by awareness, non-judgment, and acceptance of internal and external experiences. Stress is associated with numerous psychological and physical health concerns including depression and anxiety. Mindfulness-based interventions (MBIs) have consistently been shown to reduce perceived stress and improve a variety of other health outcomes. However, the mechanisms through which mindfulness confers these benefits are not yet fully understood. This study aimed to examine (1) the relationship between trait mindfulness and changes in cortisol following a negative Mood Induction Procedure (MIP) and (2) whether trait mindfulness moderates the relationship between changes in cortisol and mood reactivity following a negative MIP. Participants were recruited from the Dallas-Fort Worth Metroplex (N = 25 participants) and completed a series of self-report questionnaires, negative and positive mood induction, and mood assessments. Participants provided salivary cortisol samples before and after negative MIPs. Results indicated that, following the negative MIP, sadness and anxiety were associated with greater reductions in cortisol. Trait mindfulness moderated these associations. Overall, these findings contribute to a broader understanding of emotional responses to negatively-valenced stressors and how they are mitigated by mindfulness. They suggest support for future applications of mindfulness-based interventions in high-stress populations.

## Introduction

Stress is often associated with a variety of psychological and physical health consequences, such as migraines [1–2], depression [3], anxiety [4], poorer sleep [5], and changes in eating habits [6]. Therefore, identifying accessible and effective strategies for regulating stress is imperative. Mindfulness-based interventions (MBIs) have emerged as a promising approach, with results suggesting that MBIs reduce perceived stress and improve well-being [7–8]. Despite these benefits, the mechanisms underlying these effects remain unclear, particularly regarding how mindfulness may influence physiological stress responses, such as cortisol, and their relationship with psychological outcomes.

### Mindfulness and Stress

Stress is a response to physical (e.g., extreme heat or cold, acute or chronic pain), psychological (e.g., public speaking, workplace demands, or oral exams), or emotional (e.g., interpersonal conflict, loss of a loved one, traumatic or intense experiences) demands. Stressors can vary in intensity, ranging from minor daily annoyances (e.g., traffic) to major life-change events (e.g., the COVID-19 pandemic), and can include both positive (e.g., starting a new job) and negative (e.g., multiple upcoming deadlines) stimuli [9–11]. Individual differences, including personality and cognitive abilities such as mindfulness, can influence how individuals interpret and respond to potential stressors [12–13].

Mindfulness is posited to reduce stress by fostering greater acceptance and awareness thereby enhancing stress-management and coping skills [14–15]. Creswell and Lindsay [16] proposed a “mindfulness-stress buffering account” in which open-curiosity, nonjudgmental awareness and acceptance toward present-moment experience influence appraisal processes and buffer stress by reducing emotional reactivity. This process is thought to promote more adaptive emotion-regulation (ER) that enhances positive coping and resilience to stress. Such an adaptive ER process gained from mindfulness, as suggested by Creswell and Lindsay [16], may buffer against stress-related symptoms (e.g., migraines, chronic pain, inflammation response) [17–19], improve quality of life [19–21], improve sleep, eating, and exercise habits, and reduce deleterious stress-habits such as smoking or drug use [22–28].

### Cortisol Responses to Emotions and Stress

Cortisol is a commonly used physiological marker of stress. When under stress, the body activates “fight-or-flight” responses to manage perceived threats or negative emotions such as anxiety [29]. The sympathetic-adrenal-medullary (SAM) and the hypothalamic-pituitary-adrenal (HPA) axes underlie these responses by several mechanisms including the release of hormones like cortisol [29]. Cortisol helps sustain the energy “boost” (i.e., adrenaline) needed to respond to the stressor [29]. However, overproduction or underproduction of cortisol can impair ER of stress-concomitant feelings of anxiety, depression, and negative mood (e.g., sadness) [29]. While research consistently shows that mindfulness reduces perceived stress [30], the physiological mechanisms by which mindfulness mitigates stress-responses remain unknown. It might be that mindfulness promotes ER thereby reducing perceived stress and the ensuing cortisol response.

Existing studies on mindfulness and cortisol have produced inconsistent findings. Mindfulness training studies consistently report mood improvements [31], however some studies report concomitant increases [32], whereas others report decreases [33–37], or no change [38–43] in cortisol levels following mindfulness training. Inconsistencies in cortisol reactivity following mindfulness training may result from between-study differences in the length of training (e.g., 8-weeks vs 5 days), differences in the inclusion of, and types of stressors (e.g., mood induction, no induction, Trier Social Stress Test; TSST), and time of day (e.g., morning or late afternoon).

It is important to recognize that relationships between stress and cortisol reactivity are not straightforward. Cortisol is important in maintaining physiological and emotional function. Extremely small responses or large changes in cortisol may be indicative of an insufficient physiological stress response in the body [29]. Blunted or inadequate responses to stress are associated with adverse psychological outcomes such as depression and anxiety [3]. However, elevated or exaggerated responses are also associated with other adverse health outcomes (e.g., heart disease) [3]. Therefore, more research is needed to understand physiological stress response mechanisms, how they are indexed by cortisol and how they might be mediated by stress-related mood changes.

Despite extensive research on cortisol in stress, there is a limited understanding of how cortisol responds to other negative moods, particularly sadness [34, 44–49]. Velten Mood Induction Procedures (VMIP) consistently induce depressed mood [50–51] but have produced heterogenous results with some studies reporting increases [52–54] and others decreases [34, 45] in cortisol reactivity. Indeed, the decoupling between stress, emotions, and cortisol responses remain complex and poorly understood [49]. Furthermore, the mechanisms by which mindfulness may influence these relationships remain unclear.

Together, these findings suggest that changes in cortisol should be interpreted in the context of subjective mood, as physiological and psychological responses may diverge (i.e., a loose coupling of physiological–psychological responses) during emotion regulation [49, 55–56]. One factor that may contribute to inconsistent stress- and emotion-cortisol findings is individual differences in emotion regulation, influenced by trait mindfulness. Mindfulness is known to mitigate subjective emotional reactivity to negative mood induction procedures (MIPs) [34, 51, 57–58]. Therefore, the present study aims to assess the relationships between cortisol, trait mindfulness, and mood. Specifically, we sought to ascertain whether trait mindfulness would moderate relationships between cortisol and mood following negative MIP.

## Materials and Methods

Twenty-five participants were recruited from the Dallas-Fort Worth Metroplex through flyers, word-of-mouth, and The University of Texas at Dallas SONA participant pool. This study was part of a larger 8-week study examining the benefits and effects of a mindfulness training intervention. The recruitment period for this study was December 19, 2024, to October 3, 2025. All participants provided written informed consent before participating in the study. Participants were excluded during recruitment if they were younger than 19 years old, pregnant, had a diagnosis of color blindness, history or diagnosis of mental health disorders, were currently taking anti-depressants, psychotropic medication (excluding anxiolytics), and/or other medications for mental health disorders or illnesses, currently taking oral contraceptives, estrogens, anti-inflammatory medications (e.g., antihistamines/allergy medications, asthma medications), had oral injuries or inflammation, or history of suicide attempts or suicidal ideation. They were also excluded if they had any regular formal mindfulness-meditation practice. Formal practice was defined as regular practice of mindfulness-meditation of at least one hour per week for three months or having a total lifetime meditation practice of 12 hours. Proficiency in English was required because the negative MIP and meditation intervention required listening to, reading, and responding to prompts in English. Participants received either up to 4 research course credits or up to $100 in the form of a gift card upon full completion.

### Demographics

Participants’ ages ranged from 19 to 81 years old (6 males, 19 females). The racial composition of the sample was 40% White/Caucasian, 36% Asian, 12% Black/African American, and 12% as other. Sixteen percent reported identifying as Hispanic or Latino, and 83.33% of participants reported English as their primary language, and 16.67% reported English as a second language. Additionally, 45.83% of participants reported being bilingual and 16.67% reported knowing more than 2 languages.

### Procedure

Due to the nature of the study, suicidal ideation was assessed via Item 9 on Beck’s Depression Inventory was administered during recruitment and prior to beginning the study procedures. If any value other than 0 (“I don’t have any thoughts of killing myself”) was given, the participant was disqualified. During the in-person sessions, Beck’s Depression Inventory was re-administered before the negative MIP to screen suicidal ideation using Item 9. Two participants endorsed a value of 1 (“I have thoughts of killing myself, but I would not carry this out”), for these participants the researcher followed a suicidal ideation safety protocol prescribed by the IRB. In this protocol, the experimenter confirmed that the participant was aware of the potentially triggering stimuli in the MIP and provided contact information for the university counseling center and the hotline crisis center. No participants endorsed a value of 2 or higher (“I would like to kill myself” or “I would kill myself if I had the chance”) on BDI-II Item 9.

All testing sessions occurred between 12 noon and 6 PM to control for diurnal cortisol fluctuations. As can be seen in Fig 1, informed consent was obtained from all participants, followed by demographics. The demographic questionnaire asked about age, gender, race, ethnicity, highest education achieved, native language, and additional languages spoken. During the in-person sessions, participants completed self-report questionnaires including the Five Facet Mindfulness Questionnaire (FFMQ), Beck Depression Inventory (BDI-II), among others that were completed as part of the larger training study.

**Fig 1.**
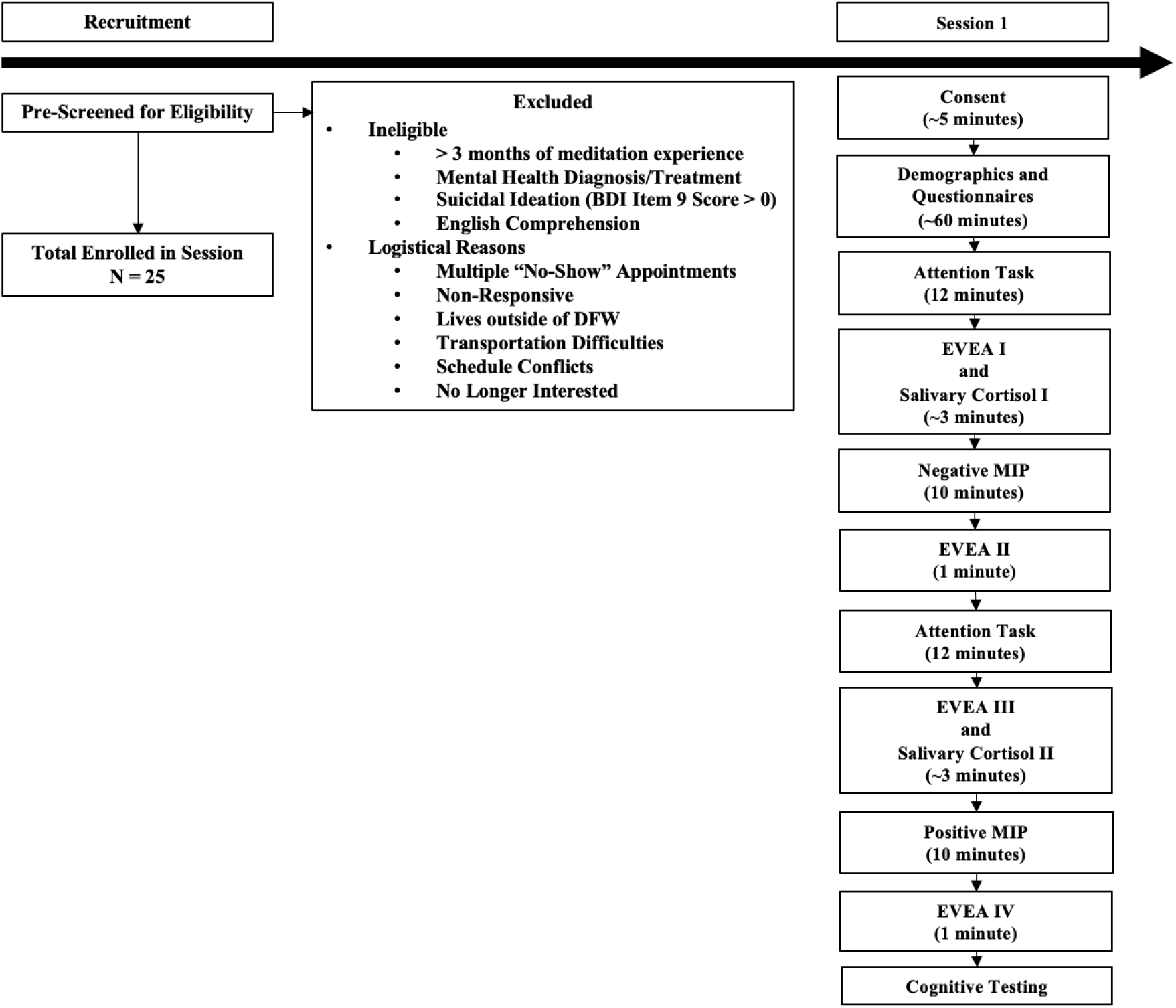
Study flow chart depicting participant recruitment, screening, and session one procedures for the 8-week Focused Attention Mindfulness Training and Relaxation Training study. The present study was conducted as part of a larger 8-week study examining the benefits and effects of mindfulness training. The present analyses focused on data collected during session 1. EVEA = Scale for Mood Assessment (Escala de Valoración del Estado de Ánimo); MIP = Mood Induction Procedure.

Following the questionnaires (Fig 1), an attention task (part of the larger study) was administered followed by the first (baseline mood) Scale for Mood Assessment (Escala de Valoración del Estado de Ánimo; EVEA). After arrival, completion of self-questionnaires, and baseline attention task (approximately 50 to 80 minutes), the first (baseline) saliva sample was taken. Participants were instructed to place a swab under their tongue for two minutes, while avoiding contact with teeth and refraining from chewing on the swab. After removing the swab, the negative MIP was administered. During this time, the researcher stepped out of the room to allow participants to complete the task privately and reduce potential social pressure of suppressing emotional responses. Immediately following the negative MIP, participants completed a second EVEA and were re-administered the attention task. A third EVEA measure was collected after the attention task. A second cortisol sample was collected about 25 minutes after completion of the attention task. Following the second cortisol sample, a positive MIP was administered to negate the effects of the negative MIP, followed by a final EVEA. The researcher then administered a final set of cognitive tests. To conclude the in-person session, all participants were debriefed with materials on information for the university counseling center and National Suicide Prevention Lifeline and the National Hopeline Network.

### Questionnaires

#### Five Facet Mindfulness Questionnaire

The Five Facet Mindfulness Questionnaire (FFMQ) [59] measures overall trait mindfulness and its five facets. These include: 1) Observe, noticing internal and external events, emotions, or thoughts (e.g., “I pay attention to how I feel at the moment in considerable detail.”), 2) Describe, defining sensations, thoughts, or feelings (e.g., “Even when I’m feeling terribly upset, I can find a way to put it into words.), 3) Act with awareness, focusing on the present-moment (e.g., “I find it difficult to stay focused on what’s happening in the present.”), 4) Nonjudgement, being non-critical of oneself (e.g., “I tell myself I shouldn’t be feeling the way I’m feeling.”), and 5) Nonreactive, allowing present thoughts or feelings to not affect the individual (e.g., “In difficult situations, I can pause without immediately reacting.). 19 of 39-items were reversed scored (e.g., “I find myself doing things without paying attention.”). Nonreactivity had 7-items while all other facets had 8-items each.

Participants responded on a 5-point Likert scale (from 1 = never or very rarely true to 5 = very often or always true). Scores were calculated by summing each subscale independently and by summing a total score for overall trait mindfulness. Higher scores indicated greater trait mindfulness or for each facet of mindfulness.

#### Scale for Mood Assessment (Escala de Valoración del Estado de Ánimo)

The Scale for Mood Assessment (Escala de Valoración del Estado de Ánimo (EVEA)) [60] was used to assess current mood. The EVEA consists of 16 items measuring four subscales of emotions, each containing 4-items: 1) Sadness-Depression (“I feel melancholy”, “I feel depressed”, “I feel downcast”, “I feel sad”), 2) Anxiety (“I feel nervous”, “I feel tense”, “I feel anxious”, “I feel restless”), 3) Anger-Hostility (“I feel irritated”, “I feel angry”, “I feel annoyed”, I feel displeased”), and 4) Happiness (“I feel happy”, “I feel optimistic”, “I feel joyful”, “I feel cheerful”). Participants responded on a 10-item Likert scale (0 = Not at all to 10 = Very Much). Scores were calculated by averaging each subscale. Mood reactivity was calculated as the difference (after – before) between participants’ baseline EVEA and their EVEA following negative mood induction. Additionally, a composite Sadness-Anxiety score was constructed *a priori* (by summing Sadness-Depression and Anxiety ratings), as an indicator of affective distress because sadness and anxiety are commonly conceptualized as internalizing forms of distress and frequently co-occur as a disorder [61–64]. In contrast, anger reflects a distinct externalizing form of distress [62, 65] and was therefore retained as a separate affective dimension, consistent with previous literature [66]. Happiness reflects positive affect rather than affective distress [67–68] and was therefore also retained as a separate affective dimension. Rumination has been posited to moderate the relationship between sadness-anxiety [69]. Therefore, sadness-related affect was included to capture sadness related affect from the negative MIP [51], while anxiety was included to capture stress-related affective responses.

#### Mood Induction Procedures (MIP)

The mood induction procedures were drawn from an established laboratory protocol [51]. Prior to mood induction procedures, participants were briefly presented with two classical music excerpts (Barber, 1939, Adagio for Strings, op. 11 [70]; Albinoni, 1981, Adagio in G Minor for organ and strings [71]). Each was played for 15 seconds, after which participants selected the piece they perceived as more “somber,” “low,” or “down.” To reduce demand characteristics, care was taken to ensure that the words “negative,” “sad,” or “depressing” were avoided throughout the procedure. Following music selection, participants listened (via E-Prime software version 2.0.10; Dell 15 inch monitor) to the music while reading statements ranging from neutral (e.g., “Today is neither better nor worse than any other day”) to increasingly negatively-valenced (e.g., “I want to go to sleep and never wake up”). They were instructed to reflect on personal experiences consistent with each statement and press the “1” or “2” key on the keyboard to indicate whether they remembered a time in their life described by the statement (key “1”) or not (key “2”). Text was displayed in black on a dark gray background. The task was self-paced and typically required approximately 10 minutes.

Following all procedures, participants underwent a positive mood induction to restore mood to baseline. The positive MIP mirrored the structure of the negative condition but used more upbeat musical selections (Beethoven, 1805, Piano Concerto No. 4, Op. 58 in G Major: III. Rondo, Vivace [72]; Mozart, 1787, Serenade No. 13 KV 525 G Major: I. Serenade. Allegro [73]). Participants selected the piece they perceived as more “upbeat”. Again, emotional descriptors such as “happy” or “positive” were avoided. Participants completed another 60 Velten statements [74], now progressing from neutral (e.g., “Today is neither better nor worse than any other day”) to increasingly positively valenced (e.g., “Wow! I feel great!”), while recalling corresponding experiences. The statements were displayed in yellow text on a bright blue background, and the task duration was approximately 10 minutes.

#### Salivary Cortisol Collection

Two saliva samples were collected using a SalivaBio Oral (Salimetrics SalivaLab; Carlsbad, CA). Participants were instructed to avoid the following prior to collection: 1) consuming foods high in sugar or acidity the day of their appointment, 2) consuming alcohol, caffeine, nicotine, or prescriptions/over-the-counter medicines within 12 hours of their appointment, 3) engaging in vigorous exercise within 12 hours of their appointment, 4) having any oral injuries or inflammations, 5) eating a major meal within two hours of their appointment, 6) consuming dairy products, 7) using hormone-based oral contraceptives or estrogen treatments, and 8) taking steroid-based anti-inflammatory medications or other medicines known to alter cortisol levels.

Prior to assay, samples were stored at -80°C before being shipped on dry ice to the Salimetrics SalivaLab (Carlsbad, CA). Samples were thawed to room temperature, vortexed, and then centrifuged for 15 minutes at approximately 3,500 RPM (1,500 x g) immediately before performing the assay. Samples were tested for salivary cortisol using a high sensitivity enzyme immunoassay (Cat. No. 1-3002). The sample test volume was 25 μl of saliva per determination. The assay has a lower limit of sensitivity of 0.007 μg/dL, a standard curve range from 0.012 -3.0 μg/dL, and an average intra-assay coefficient of variation of 4.60%, and an average inter-assay coefficient of variation of 6.00%, which meets the manufacturers’ criteria for accuracy and repeatability in Salivary Bioscience, and exceeds the applicable NIH guidelines for Enhancing Reproducibility through Rigor and Transparency. Samples were assayed at the Salimetrics’ SalivaLab (Carlsbad, CA) using the Salimetrics Salivary Cortisol Assay Kit (Cat. No. 1-3002), without modifications to the manufacturers’ protocol.

### Statistical Analysis

All statistical analyses were conducted in R [75] using RStudio [76]. Descriptive analyses evaluated internal consistency and reliability (using Cronbach’s alpha), influential data points (using Cook’s distance), skewness (using Shapiro-Wilk test). Differences in cortisol and mood from baseline to post-negative mood induction was assessed using paired-sample *t*-tests). Additionally, an *a priori* power analysis was conducted using G*Power 3.1 [77] to estimate the necessary sample size based on effect size calculated from the reported means and standard deviations of cortisol responses following a negative mood induction in a comparable experimental paradigm [44]. Primary analyses included testing associations between mood reactivity, changes in cortisol, and trait mindfulness (using simple linear regressions). Changes in cortisol and mood reactivity were calculated as differences in post-induction to baseline scores. Participants were dummy coded into two groups based on reactivity to the negative mood induction (0 = non-reactive; 1 = reactive). A moderation analysis was then performed to assess influence of trait mindfulness on mood reactivity-based group differences in cortisol change.

### Power Analysis

An *a priori power analysis* was conducted using G*Power 3.1 [77]. Parameter estimations were based on an effect size (d = 0.81) calculated from the reported means and standard deviations of salivary cortisol responses following a negative mood induction in a comparable experimental paradigm [44]. Assuming an alpha level of .05 and statistical power of .80, the analysis indicated that a minimum sample of 14 participants was required to detect the expected effect for a paired-samples comparison of cortisol responses.

## Results

### Descriptive Statistics

Cronbach’s alpha, calculated for the overall FFMQ, was found to be 0.93 indicating excellent internal consistency and reliability. Consistency for the individual facets were as follows: observe = 0.70, describe = 0.92, act with awareness = 0.69, nonjudgement = 0.93, and nonreactive = 0.69. These values indicated acceptable to excellent reliability across all facets. EVEA at baseline showed excellent reliability (Cronbach’s alpha = 0.92). Excellent consistency was observed for the individual mood subscales, as follows: sadness-depression = 0.93, anxiety = 0.85, sadness-anxiety = 0.93, anger-hostility = 0.90, happiness = 0.95. After negative MIP, EVEA still displayed excellent consistency (Cronbach’s alpha = 0.93). Excellent consistency continued to be observed for the individual mood subscales as follows: sadness-depression = 0.95, anxiety = 0.89, sadness-anxiety = 0.95, anger-hostility = 0.89, happiness = 0.93. For the remaining participants (N = 23), Table 1 provides descriptive statistics (means and standard errors) for trait mindfulness, cortisol and mood. Participants reported an average trait mindfulness score of 122.63 on the FFMQ.

Cook’s distance analysis was conducted to assess influential data points. Two participants’ cortisol change values exceeded four standard deviations from the sample mean and demonstrated high influence in the Cook’s distance analysis. These two participants were excluded from further analyses. Raw baseline cortisol levels were observed to be skewed (skewness = +2.27). Raw changes in cortisol levels were also skewed (skewness = -3.01). Therefore, raw cortisol values were log-transformed. Change was calculated after log-transformation. A Shapiro-Wilk test confirmed normality after log-transformation for changes in cortisol (W = 0.94, p = 0.20), baseline cortisol (W = 0.98, p = 0.95), and after MIP cortisol (W = 0.94, p = 0.53).

**Table 1:**
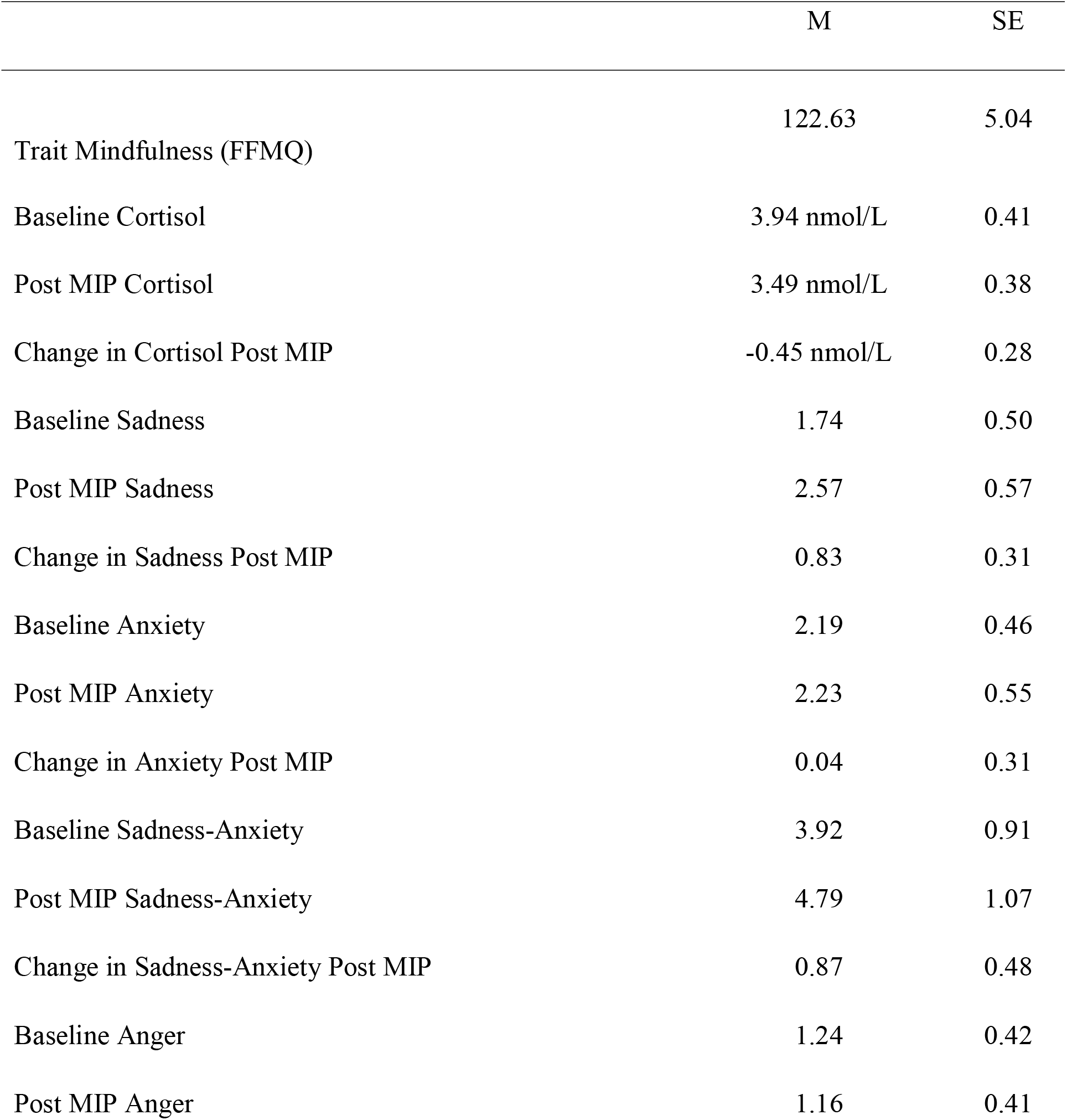
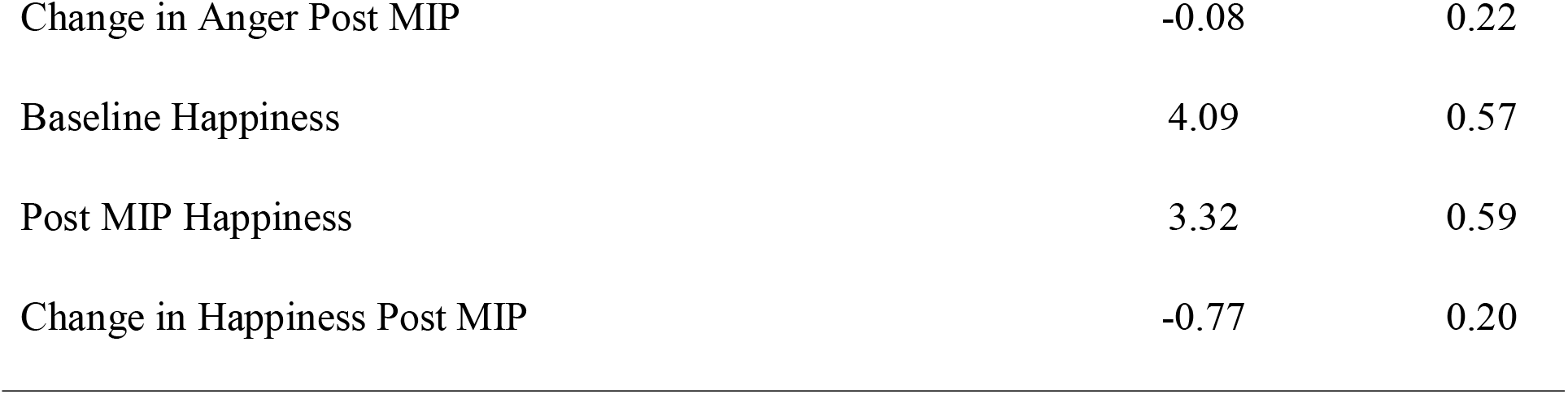
Descriptives (Means and Standard Errors) for Trait Mindfulness and Changes in Cortisol and Mood Following MIP.

|  | M | SE |
|---|---|---|
| Trait Mindfulness (FFMQ) | 122.63 | 5.04 |
| Baseline Cortisol | 3.94 nmol/L | 0.41 |
| Post MIP Cortisol | 3.49 nmol/L | 0.38 |
| Change in Cortisol Post MIP | -0.45 nmol/L | 0.28 |
| Baseline Sadness | 1.74 | 0.50 |
| Post MIP Sadness | 2.57 | 0.57 |
| Change in Sadness Post MIP | 0.83 | 0.31 |
| Baseline Anxiety | 2.19 | 0.46 |
| Post MIP Anxiety | 2.23 | 0.55 |
| Change in Anxiety Post MIP | 0.04 | 0.31 |
| Baseline Sadness-Anxiety | 3.92 | 0.91 |
| Post MIP Sadness-Anxiety | 4.79 | 1.07 |
| Change in Sadness-Anxiety Post MIP | 0.87 | 0.48 |
| Baseline Anger | 1.24 | 0.42 |
| Post MIP Anger | 1.16 | 0.41 |
| Change in Anger Post MIP | -0.08 | 0.22 |
| Baseline Happiness | 4.09 | 0.57 |
| Post MIP Happiness | 3.32 | 0.59 |
| Change in Happiness Post MIP | -0.77 | 0.20 |

### Mood and Cortisol Manipulation Check

There were significant decreases in cortisol levels following the negative MIP. A paired sample t-test indicated a significant decrease in cortisol values following the negative MIP, ΔM = -0.06, 95% CI [-0.12, -0.004], *t*(22) = -2.22, *p* = 0.04 (Cohen’s d = -0.46; see Fig 2). The paired-sample t-test indicated a marginal significance for increases in sadness-anxiety following negative MIP, ΔM = 0.87, 95% CI [-1.87, 0.13], *t*(22) = 1.81, *p* = 0.08, with a moderate effect size (Cohen’s d = 0.38; see Fig 3). The paired-sample t-test indicated a significant increase in sadness following negative MIP, ΔM = 0.83, 95% CI [0.11, 0.99], *t*(22) = 2.67, *p* = 0.01, with a moderate effect size (Cohen’s d = 0.56; see Fig 3). The paired sample t-test indicated a significant decrease in happiness following negative MIP, ΔM = -0.77, 95% CI [-0.32, -1.26], *t*(22) = -3.82, *p* = 0.0009 with an effect size (Cohen’s d = -0.80). However, paired-sample t-tests indicated that anger and anxiety were not significantly different following negative MIP, (p > .05).

**Fig 2.**
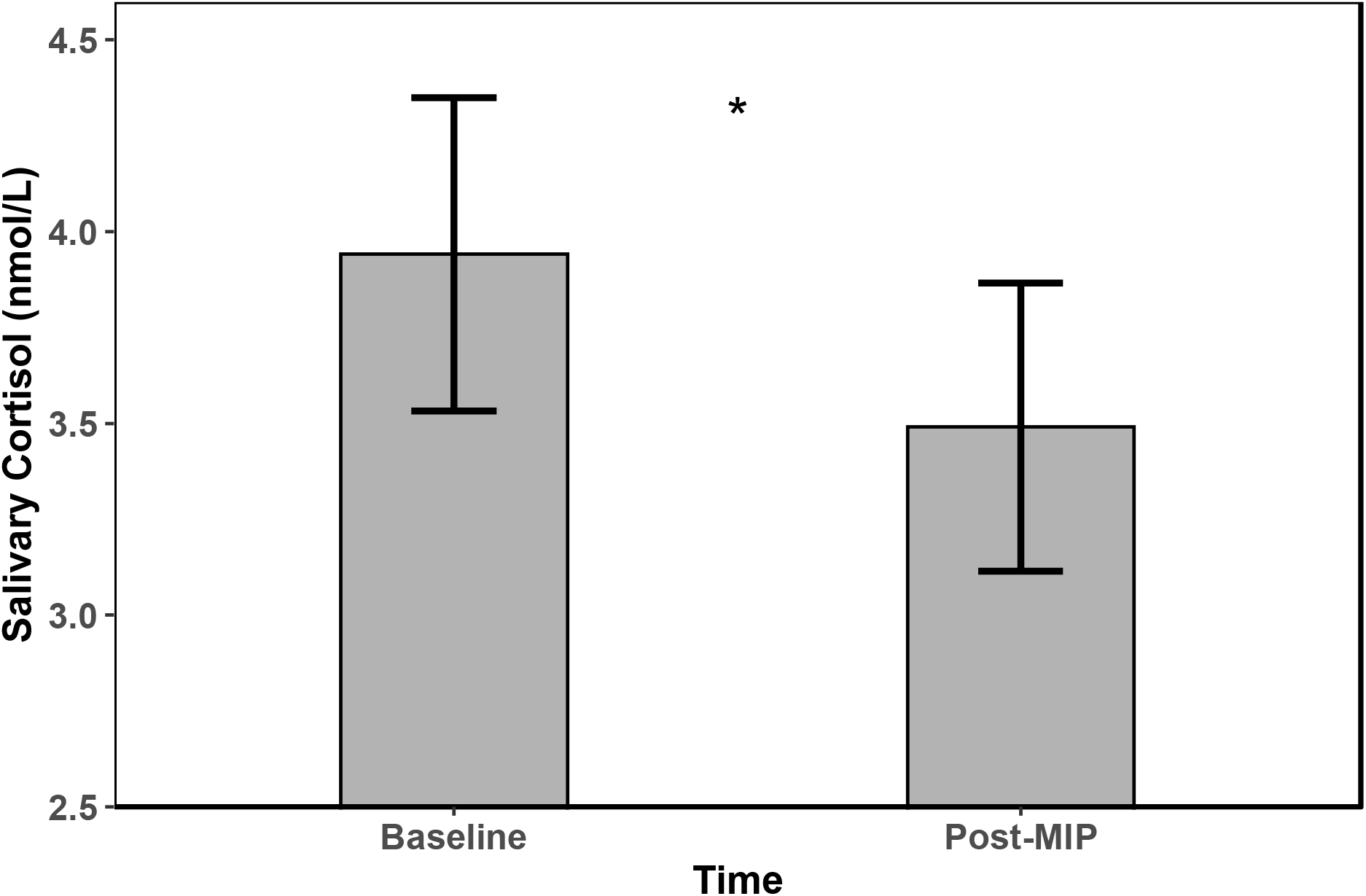
Salivary Cortisol (log nmol/L) was measured at baseline and post negative MIP. Error bars represent ±1 standard error of the mean. For visualization purposes, raw cortisol values were used in the figure.

**Fig 3.**
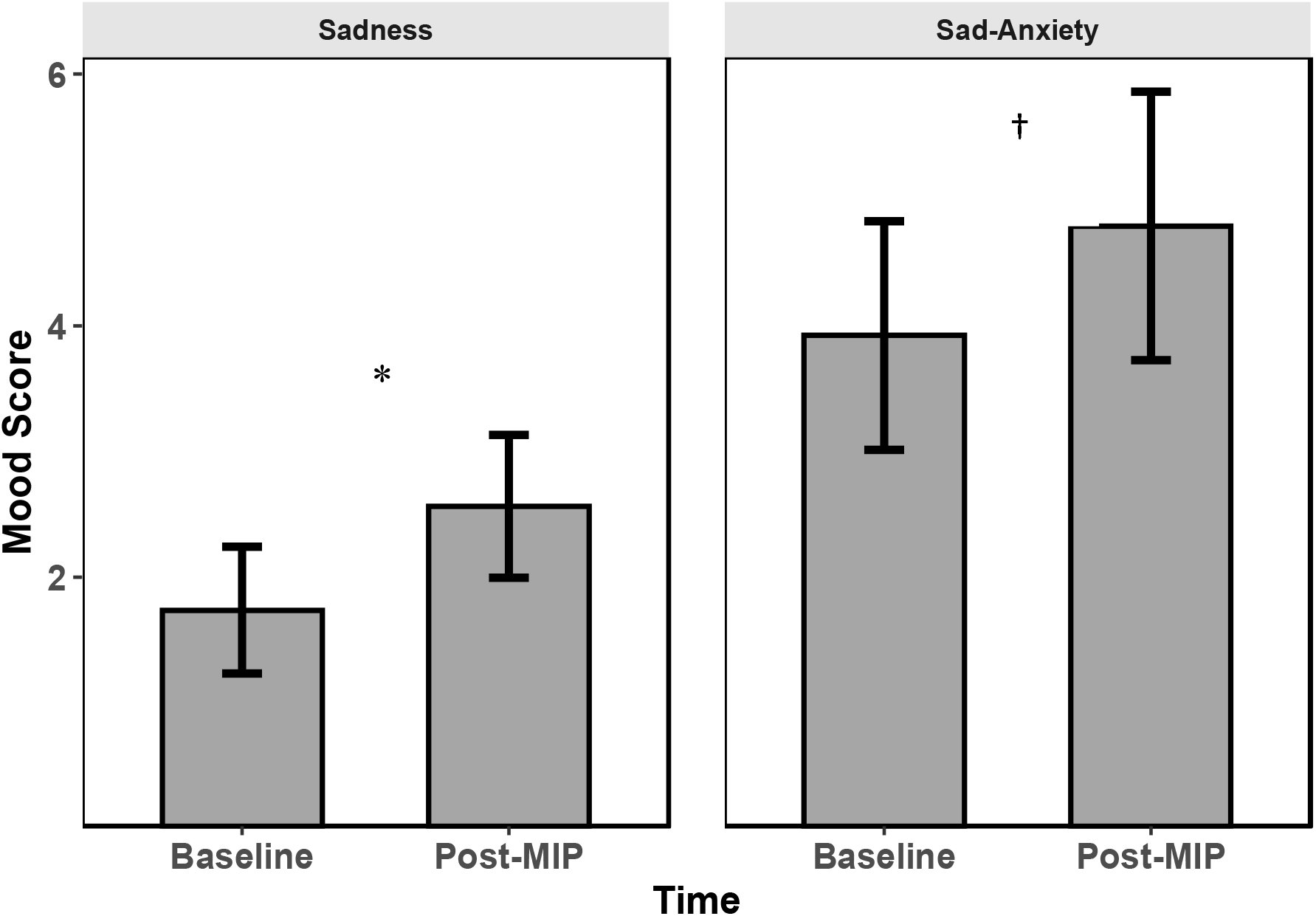
EVEA scores for sadness and sadness-anxiety were measured at baseline and post negative MIP for the cortisol subsample. Error bars represent ±1 standard error of the mean. Asterisk indicates significant mood change for sadness (p = .01) and a dagger indicates a marginal significance for sadness-anxiety (p=0.08).

### Primary Analyses

#### Regression Analyses

Simple linear regression models were conducted to examine associations between sad and sadness-anxiety mood reactivity, changes in cortisol and trait mindfulness (see Table 2). Greater decreases in cortisol were significantly associated with greater increases in sadness-anxiety following the negative MIP, while changes in cortisol were not significantly associated with sadness. Higher trait mindfulness was associated with reduced sadness and sadness-anxiety reactivity, whereas trait mindfulness was not associated with changes in cortisol following negative MIP (see Fig 3).

**Table 2:** Regressions among Trait Mindfulness and Changes in Cortisol and Mood Following MIP.

|  |  | β | *F(1,21)* | *p* | R<sup>2</sup> |
|---|---|---|---|---|---|
| Δ Cortisol | Δ Sadness | -0.361 | 3.146 | 0.09 | 0.130 |
| Δ Cortisol | Δ Sadness-Anxiety | -0.577 | 10.47 | 0.004 | 0.333 |
| Δ Cortisol | Trait Mindfulness (FFMQ) | -0.040 | 0.033 | 0.858 | 0.002 |
| Δ Sadness | Trait Mindfulness (FFMQ) | -0.542 | 8.744 | 0.008 | 0.294 |
| Δ Sadness-Anxiety | Trait Mindfulness (FFMQ) | -0.450 | 5.337 | 0.031 | 0.203 |

**Fig 4.**
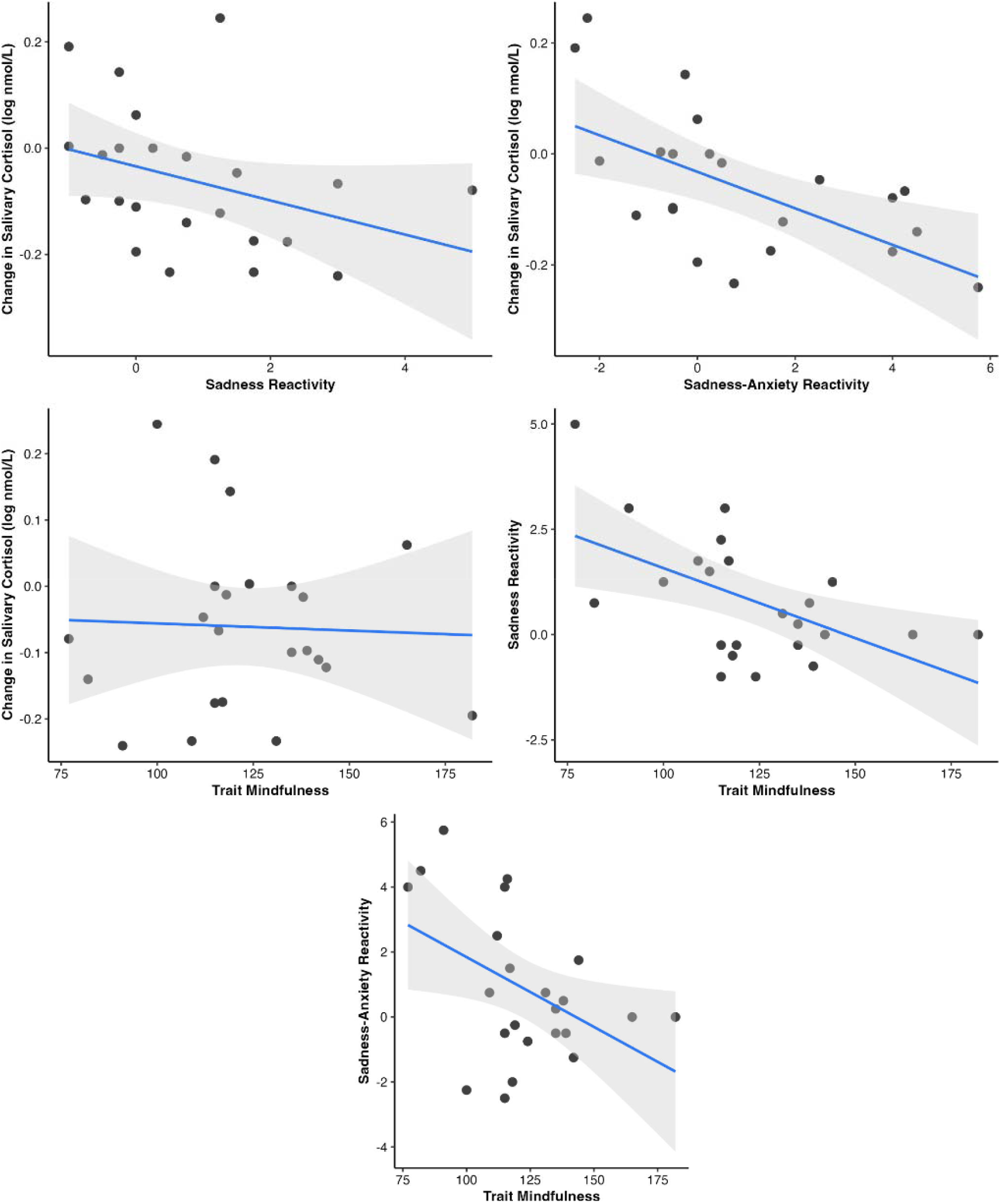
Associations between trait mindfulness, cortisol, and mood reactivity following the negative mood induction. Composite sadness-anxiety scores were calculated by summing sadness and anxiety and computing the change from baseline to post-induction.

#### Trait Mindfulness and Psychophysiological Reactivity

To account for individual differences in responses to the negative mood induction (Clark, 1983), participants were grouped according to those whose mood worsened (reactors) versus those whose mood improved or did not change (non-reactors) in response to the negative MIP, to account for individual variability in mood-induction responses [78]. We performed a moderation analysis to assess trait mindfulness and mood reactivity influences on changes in cortisol. Trait mindfulness significantly moderated the association between sadness-anxiety reactivity and cortisol change, (b = 0.005, SE = 0.002, p = 0.015). The overall model was significant, F(3, 19) = 7.35, p = 0.002, accounting for 54% of the variance in cortisol change. Low and high trait mindfulness were operationalized as 1 standard deviation (SD) below and above the sample mean, respectively. Simple slopes analyses revealed that, among low trait mindfulness individuals, a significant interaction between sadness-anxiety reactivity was associated with decreases in cortisol after negative MIP, b = -0.29, SE = 0.06, p < 0.001; however, for high trait mindful individuals, no significant moderation effects were observed for mood reactivity and changes in cortisol after negative MIP, b = -0.05, SE = 0.06, p = 0.45. Thus, higher trait mindfulness attenuated the relationship between sadness-anxiety reactivity and changes in cortisol. No such effects were observed when sadness and anxiety was used as the moderator. Descriptive statistics for reactors versus non-reactors after negative MIP are provided in Table 3. Therefore, high trait mindfulness attenuated the relationship between sadness-anxiety reactivity and changes in cortisol (Fig 5).

**Table 3:**
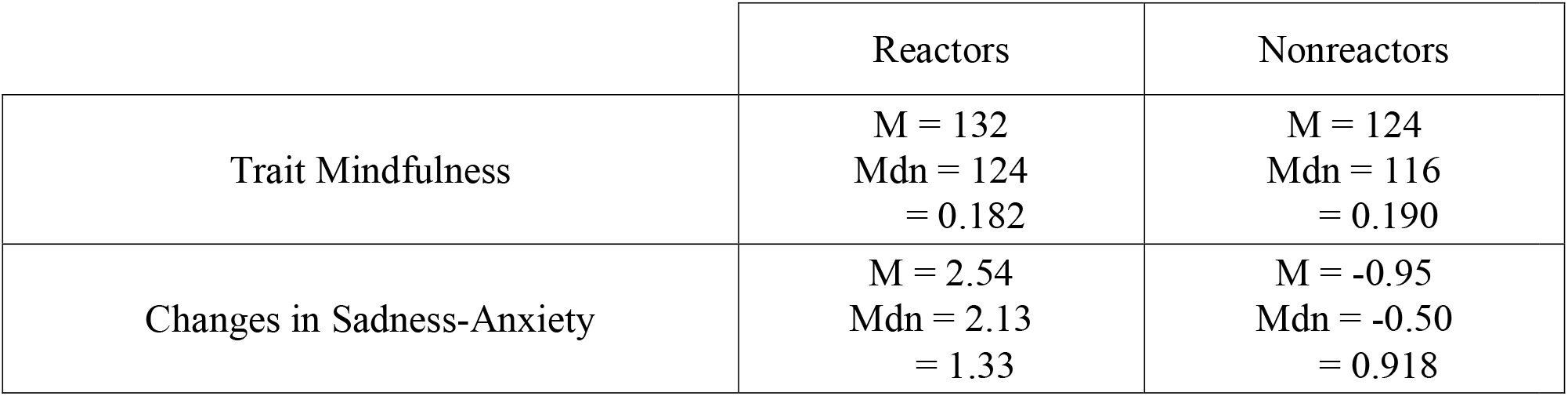
Descriptives (Means, Median, and Coefficient of Variance) for Trait Mindfulness and Changes in Sadness-Anxiety Based on Reactivity.

|  | Reactors | Nonreactors |
|---|---|---|
| Trait Mindfulness | M = 132<br>Mdn = 124<br>= 0.182 | M = 124<br>Mdn = 116<br>= 0.190 |
| Changes in Sadness-Anxiety | M = 2.54<br>Mdn = 2.13<br>= 1.33 | M = -0.95<br>Mdn = -0.50<br>= 0.918 |

**Fig 5.**
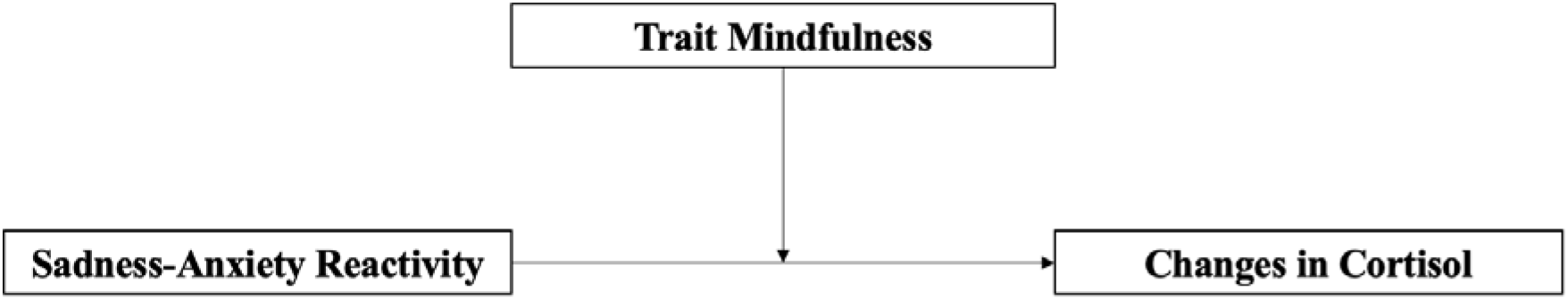
The moderation model. N = 23.

## Discussion

In this study, we examined relationships between trait mindfulness, cortisol, and mood following a negative MIP. Guided by the mindfulness stress-buffering account [16] and the transactional model of stress [12], we hypothesized that higher trait mindfulness would attenuate cortisol level changes and mood reactivity following a negative MIP. Consistent with these frameworks, the present findings suggest that trait mindfulness may function as an important individual-difference dimension in stress reactivity.

The negative MIP increased feelings of sadness-anxiety and sadness, and decreases of happiness, indicating that the procedure effectively induced negative emotions. However, cortisol levels were significantly reduced following the MIP, contrary to our initial expectation. Trait mindfulness also did not predict cortisol levels following the MIP; however, greater decreases in cortisol were associated with greater increases in sadness-anxiety and anxiety reactivity following the MIP. These findings suggest a more complex relationship between cortisol and emotional reactivity than previously understood. Cortisol reactivity following a stressor, such as the TSST, is typically characterized by increases in cortisol, reflecting activation of a “fight-or-flight” response [79–81]. However, in the context of the negative MIP, greater cortisol reductions were observed and were associated with greater increases in sadness-anxiety reactivity. These findings suggest that cortisol responses may differ depending on the emotional context. It might be that whereas social stressors elicit increases in cortisol [79–81], negative mood inductions elicit decreases in cortisol [45, 82].

In this study, trait mindfulness moderated the relationship between cortisol change and sadness-anxiety reactivity. Our findings from the moderation model for trait mindfulness and cortisol change would suggest that mindfulness may attenuate the association between physiological reactivity and emotional responses to stress. This pattern is consistent with the Mindfulness-Stress Buffer Hypothesis [16] which proposes that mindfulness attenuates the impact of stress on psychological outcomes. Consistent with this hypothesis, probing the interaction revealed that at 1 SD below the mean of trait mindful, greater sadness-anxiety reactivity was associated with greater decreases in cortisol following negative MIP. However, this association was reduced among individuals with high trait mindfulness. It might be that mindfulness weakens the link between cortisol reactivity and negative-mood reactivity, possibly by creating a buffer between changes in psychological mood and their associated physiological responses.

## Conclusions

This study was limited by a relatively small sample size. However, the present findings provide preliminary evidence for cortisol reactivity following a negative MIP, contributing to a broader understanding of acute stress-responses such as sadness-anxiety induced by MIPs. More work will be needed to compare social stressors induced by, for instance, the TSST and mood stressors such as those induced by the MIPs. Using EVEA, we observed that, whereas all participants showed equivalent EVEA scores at baseline, there was a group of individuals for whom mood changed following the MIP more than those for whom mood was equivalent before and after the MIP. Future studies will be needed to understand the precise mechanisms of these individual differences.

A further limitation is that the study was restricted to two cortisol sampling time points, which limits the ability to capture the full range of possible cortisol changes to the negative MIP. To assess cortisol change following mood manipulation, the second saliva sample was collected 25 minutes following the negative MIP, consistent with the expected peak in cortisol response to the MIP [83]. Baseline cortisol was also assessed about one hour after arrival to the laboratory to account for anticipatory stress (the so-called “white coat effect”). However, future research would benefit from including additional samples to more rigorously assess cortisol changes over time. More work is needed to understand cortisol level changes and their relationship to stress and mood [84]. Despite these limitations, the study provides novel insight into the relationship between trait mindfulness, subjective mood, and physiological responses following a negative mood induction.

## Acknowledgments

The authors wish to thank our undergraduate and graduate interns for their exceptional efforts on various aspects of the study including recruitment, tracking, preparing data for analyses, and technical support. Additionally, we wish to thank Dorsey Standish for providing support and consultation throughout the project.

